# A full-length GRAS domain protein underpins efficient shoot regeneration in pepper

**DOI:** 10.64898/2026.04.01.715768

**Authors:** Su-Jin Park, Jin Ho Yang, Hyun-Soon Kim, Hyo-Jun Lee

**Author notes:** Correspondence (H.S.K.), (H.J.L.).

## Abstract

Pepper (*Capsicum annuum* L.) is a recalcitrant species regarding shoot regeneration, a trait that serves as a major bottleneck for the application of genetic engineering tools. In this study, comparative genetic analysis between a rare high-regeneration cultivar and a common low-efficiency cultivar identified a single nucleotide polymorphism (SNP) in *PHYTOCHROME A SIGNAL TRANSDUCTION 1* (*CaPAT1*) that determines shoot regeneration efficiency. The T478C SNP in the high-efficiency cultivar converts a stop codon into an Arg codon, leading to translational read-through into the neighboring gene and forming an intact GRAS domain. This SNP-mediated formation of full-length CaPAT1 is essential for its dimerization. Notably, the overexpression of *CaPAT1^T478C^* in multiple low-efficiency cultivars, including both hot and bell peppers, significantly improved both shoot regeneration and transformation efficiency in the transformed T0 generation. These findings demonstrate the pivotal role of CaPAT1 in enhancing shoot regeneration and provide a robust strategy to overcome recalcitrance in pepper.

## Introduction

Pepper (*Capsicum annuum* L.) is a globally important crop species, valued not only for its economic contribution but also for its abundance of nutrients and bioactive compounds that benefit human health (Hernandez-Perez et al. 2020; Suica-Bunghez et al. 2020; Kothari et al. 2010). However, pepper cultivation is often constrained by its sensitivity to environmental stresses, such as temperature fluctuations, humidity, and pathogen infection, which greatly interfere with normal growth and development, leading to low yields (Ko et al. 2016; Rajametov et al. 2021). To overcome these challenges, there is a high demand for research on the genetic elements involved in plant responses to these environmental stresses for the development of new cultivars through genetic engineering. However, pepper is a representative regeneration-recalcitrant species, particularly regarding shoot regeneration. Previous studies have reported that although callus induction occurs efficiently in pepper, the subsequent shoot regeneration efficiency is extremely low. For example, the cultivar Zunla-1 shows a callus induction rate of 97%, whereas the shoot regeneration efficiency is only 2.9% (Naeem et al. 2025). Similarly, the cultivar CM334 exhibits a callus induction rate of approximately 90%, but its shoot regeneration efficiency remains as low as 1.9%. Furthermore, several pepper cultivars, including Takanotsume, Jeju Jaerae, Thai Hot, and Chili Bangi, are capable of callus induction but exhibit no shoot regeneration capacity under tissue culture conditions (Kim et al. 2023). Due to these limitations, many attempts to generate gene-edited pepper plants using the Clustered Regularly Interspaced Short Palindromic Repeats (CRISPR)/CRISPR-associated protein 9 (Cas9) system have not been successful (Park et al. 2021). Therefore, enhancing shoot regeneration efficiency is a prerequisite for employing modern plant biotechnology in pepper.

In tissue culture, excised tissues are cultured on media supplemented with specific combinations of auxins and cytokinins, which act as key regulators of cell fate reprogramming. The relative balance between these two hormones determines the developmental fate of cultured cells. When explants are placed on callus-inducing medium (CIM) containing a high concentration of auxins, somatic cells form callus tissues with pluripotent characteristics and root-like identity (Radhakrishnan et al. 2018; Bustillo-Avendaño et al. 2017; Ikeuchi et al. 2019). Subsequent transfer of the callus to shoot-inducing medium (SIM), characterized by a higher cytokinin-to-auxin ratio, promotes a transition in cellular identity toward shoot formation and ultimately leads to the establishment of a functional shoot meristem (Christianson and Warnick. 1983; Ikeuchi et al. 2019; Higuchi et al. 2004). In many tissue culture studies, culture conditions have been investigated to induce shoot regeneration by screening the types, concentrations, and combinations of phytohormones, adjuvant content, types of explants, and environmental conditions (Gubis et al. 2004; Mazumdar et al. 2010; Park et al. 2023). In pepper, specific culture conditions for certain genotypes have been identified to enhance shoot regeneration efficiency (Gammoudi et al. 2018; Kim et al. 2023; Shams et al. 2024). For example, in the pepper cultivars Zunla-1 and CM334, shoot regeneration was not observed under standard shoot induction conditions. However, supplementation of the medium with 10 mg/L AgNO_3_ and 1 nM REGENERATION FACTOR 1 partially enhanced regeneration, resulting in efficiencies of 2.9% in Zunla-1 and 1.9% in CM334 (Naeem et al. 2025). These results demonstrate that the optimization of culture conditions can partially enhance regeneration capacity in pepper. However, regeneration efficiency remains insufficient for genetic engineering and is highly dependent on genotypes, which makes it difficult to apply these methods to various pepper cultivars. In addition to low regeneration efficiency, pepper tissue culture frequently results in abnormal morphogenesis, characterized by the formation of leafy calli and non-viable shoots (Shin et al., 2026). These aberrant structures often fail to establish a functional shoot apical meristem, further limiting successful plant regeneration.

Plant cell reprogramming is regulated by a conserved genetic network that confers pluripotency and drives *de novo* organogenesis. Detailed molecular mechanisms and signaling pathways regarding these processes have been identified in *Arabidopsis thaliana*. During incubation on CIM, auxin-dependent activation of WUSCHEL-RELATED HOMEOBOX (WOX) family members, including *WOX11* and *WOX12*, initiates the transition of differentiated cells into callus cells (Liu et al. 2014; Ulmasov et al. 1999). The establishment of pluripotency further requires *PLETHORA* (*PLT*) genes. PLT3, PLT5, and PLT7 induce the expression of *PLT1* and *PLT2* to drive the acquisition of pluripotency in the callus on CIM (Kareem et al. 2015). When callus is transferred to SIM, PLT3, PLT5, and PLT7 trigger the expression of the *CUP-SHAPED COTYLEDON 2* (*CUC2*) gene to promote PIN-FORMED 1 polarization, specifying the shoot meristem initiation site (Gordon et al. 2009; Pulianmackal et al. 2014). Subsequently, the cytokinin-responsive transcription factor WUSCHEL (WUS) is activated through the type-B ARR signaling pathway, functioning as a central regulator that stabilizes the stem-cell niche and enables shoot regeneration (Lee et al. 2018; Meng et al. 2017; Zhang et al. 2017). While the key genes involved in shoot regeneration have been identified in Arabidopsis, whether these mechanisms are conserved and which molecular factors are responsible for the low efficiency of shoot regeneration in pepper remain largely unknown.

In addition to these mainstream signaling pathways, the GIBBERELLIN-INSENSITIVE, REPRESSOR OF GA1-3, and SCARECROW (SCR) (GRAS) domain transcription factors have been identified as being involved in shoot regeneration processes. Among the eight subfamilies of GRAS domain proteins, the PHYTOCHROME A SIGNAL TRANSDUCTION 1 (PAT1) subfamily consists of five members, specifically PAT1, SCR-LIKE 1 (SCL1), SCL5, SCL13, and SCL21 (Bolle et al. 2004). Among them, PAT1 and SCL21 were originally characterized as positive regulators of phytochrome A-mediated light signaling, as *pat1* and *scl21* mutants exhibited reduced photomorphogenic responses to far-red light (Torres-Galea et al. 2013). However, follow-up studies on PAT1 have revealed that it forms a heterodimer with ETHYLEN RESPONSE FACTOR 115 (ERF115) to enhance regenerative competence (Heyman et al. 2016). Simultaneous ectopic expression of *ERF115* and *PAT1* triggers neoplastic overgrowth that manifests as spontaneous callus formation. In addition, overexpression of these genes induces the expression of regeneration-related genes such as *PLT3*, *PLT5*, *PLT7*, and *WOUND INDUCED DEDIFFERENTIATION 1* (Kareem et al. 2015). Conversely, the *pat1* and *erf115 pat1* mutants showed low root tip regeneration capacity and impaired callus formation (Heyman et al. 2016; Bisht et al. 2023). Furthermore, PAT1 has been identified as a key factor for successful grafting in both Arabidopsis and *Picea abies* (Feng et al. 2024). These results suggest that PAT1 is involved in cell reprogramming, although its role in shoot regeneration has not yet been investigated.

In this study, we investigated the genetic factors regulating shoot regeneration efficiency using two pepper genotypes with contrasting regenerative capacities, identifying that the full-length CaPAT1 confers high shoot regeneration capacity. Our results demonstrated that ectopic expression of the full-length *CaPAT1* improved regeneration efficiency across all tested cultivars, indicating that this approach is broadly applicable to diverse pepper cultivars. This work not only elucidates the genetic basis for the regeneration-recalcitrant trait of pepper but also provides a robust platform for enhancing pepper genetic engineering through highly efficient shoot regeneration.

## Results

### TGN Represents a Pepper Genotype with High Shoot Regeneration Capacity

To investigate the genetic basis underlying low shoot regeneration efficiency in pepper, we evaluated the regeneration capacity of multiple genotypes under identical culture conditions. Among the tested genotypes, we identified two lines exhibiting contrasting regeneration phenotypes: one line with high regeneration efficiency and another with no detectable regeneration. The high-efficiency line corresponds to a previously reported transgenic pepper harboring the *J1-1* gene, which confers resistance to the fungal pathogen *Colletotrichum gloeosporioides* (Seo et al. 2014). Although this line was originally described as being derived from the cultivar ‘Nokgwang’, it exhibited markedly enhanced shoot regeneration compared to the non-regenerating ‘Nokgwang’ cultivar used in this study (Figure 1a,b). Hereafter, we refer to the high-regeneration line as TGN and the non-regenerating cultivar as N. In TGN, shoot regeneration frequency reached up to 15% at 10 weeks after incubation, whereas no shoot regeneration was observed in N. In addition, callus tissues derived from TGN displayed a significantly higher greening index compared to those of N, showing more than a two-fold increase (Figure 1c,d). Given that callus greening is closely associated with the acquisition of pluripotency and shoot regeneration competence (Heszky et al. 1989; Kareem et al. 2015), these results indicate that TGN possesses intrinsically high regeneration capacity relative to N

**FIGURE 1.**
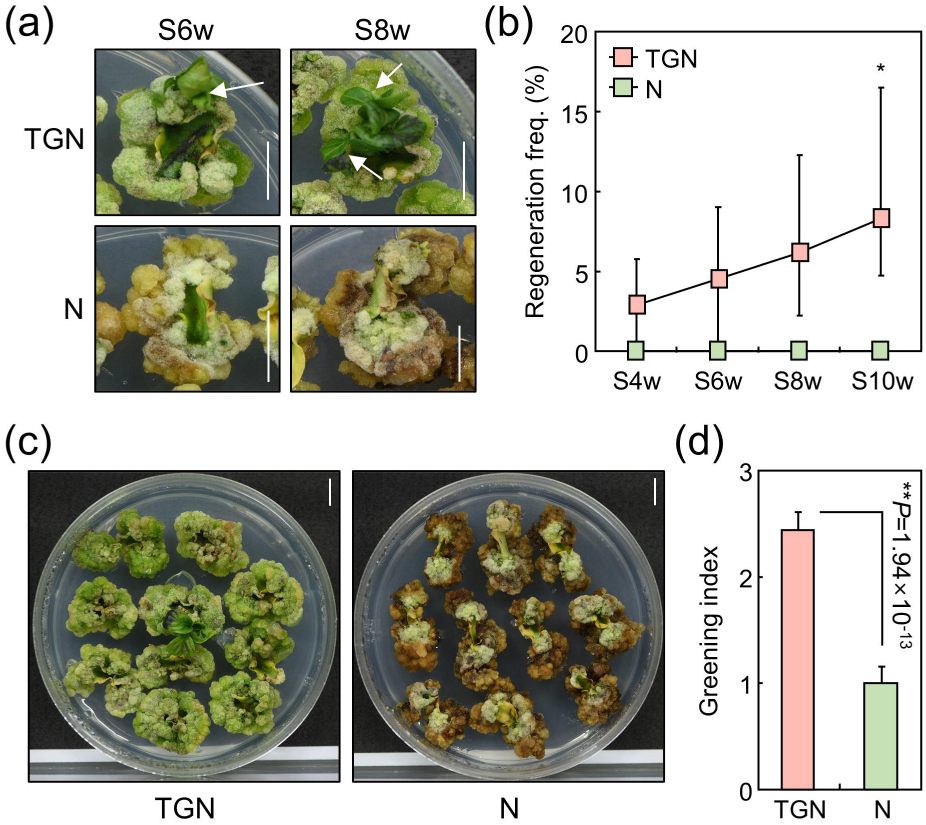
TGN exhibits high shoot regeneration capacity. (a, b) Comparing of shoot regeneration capacities between TGN and N. Cotyledon explants were utilized for the assay. Representative images showing explants developing calli and shoots are presented (a). Regeneration frequency was analyzed at the indicated time points during incubation on SIM (b). S, SIM; w, weeks. Statistical significance was determined using Student’s *t*-test (\**P* < 0.05; *n* = 3; 95 explants for each replicate). Scale bars, 1 cm. (c, d) Greening of callus in TGN and N. Representative images of explants developing callus at 8 weeks of SIM incubation are displayed (c). Greening index was quantified using ImageJ software (d). Statistical significance was determined using Student’s *t*-test (*n* = 10; 11 explants for each replicate). Scale bars, 1 cm. For line and bar graphs, whiskers indicate ± standard deviation of the mean (SD).

### Genomic Analysis Reveals That TGN and N Are Genetically Distinct Genotypes

Despite being designated under the same background cultivar name (‘Nokgwang’), we hypothesized that TGN and N are genetically distinct. This assumption was based on two observations. First, the *J1-1* gene encodes an antifungal protein with no known role in shoot regeneration. Second, the original study reporting TGN successfully achieved shoot regeneration using its background line (Seo et al. 2014), whereas N exhibited no regeneration capacity under identical conditions in our experiments. To test this hypothesis, we performed whole-genome sequencing of both TGN and N. Comparative analysis identified more than 6 million single nucleotide polymorphisms (SNPs) between the two lines (Figure 2a), clearly indicating that TGN and N represent distinct genetic backgrounds despite sharing the same cultivar name.

**FIGURE 2.**
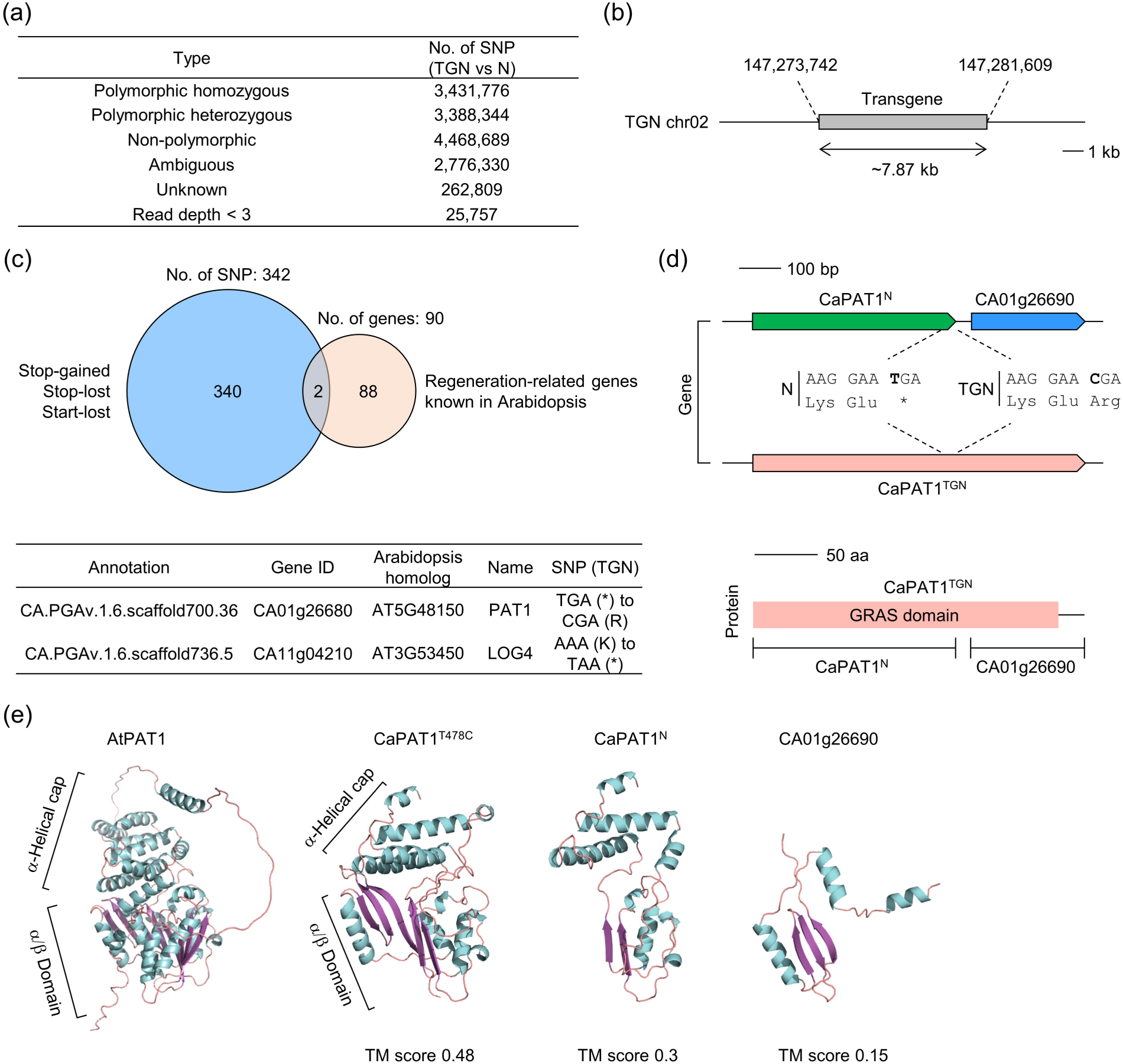
The T478C SNP produces a full-length CaPAT1 in TGN. (a) Summary of whole genome sequencing analysis in TGN and N. Sequencing reads were mapped to the pepper reference genome (Annuum.v1.6), and the total number of SNPs between TGN and N was annotated. (b) Genomic location of the transgene in TGN. The exact transgene insertion site was identified using read-split analysis and *de novo* genome assembly. Note that no endogenous genes were identified within a 3 kb range upstream or downstream of the insertion site. (c) Identification of regeneration-related genes harboring critical SNPs. Arabidopsis homologs of pepper genes containing stop-gained, stop-lost, or start-lost SNPs in TGN were compared with previously reported regeneration-related genes. The reference genome annotations for two candidate genes are listed. Gene IDs were retrieved from the CM334 genome database (https://solgenomics.net). (d) Schematic representation of the *CaPAT1* locus in TGN and N. The site of the stop-to-Arg transition in TGN is indicated. Protein domain analysis was performed using the NCBI CD-Search tool (https://www.ncbi.nlm.nih.gov/Structure/cdd/wrpsb.cgi). (e) Structural similarity analysis of Arabidopsis and pepper PAT1 proteins. Protein structures were predicted using AlphaFold3. CaPAT1^T478C^ represents the fusion protein of CaPAT1^N^ and CA01g26990 resulting from the T478C stop-lost SNP. Structural similarity was quantified by calculating TM scores using the TM-align tool (https://zhanggroup.org/TM-align/).

To exclude the possibility that the enhanced regeneration phenotype of TGN is caused by transgene insertion, we analyzed the genomic location of the transgene using whole-genome sequencing data. The analysis revealed a single-copy insertion of approximately 7.87 kb located between 147,273,742 and 147,281,609 bp on chromosome 2. This region corresponds to an intergenic region with no annotated genes within 3 kb upstream or downstream of the insertion site (Figure 2b). These results strongly suggest that the high shoot regeneration efficiency observed in TGN is unlikely to be attributable to positional effects of the transgene, and instead reflects inherent genetic differences between TGN and N.

Based on the whole genome sequencing analysis, we aimed to identify key SNPs conferring enhanced shoot regeneration capacity in TGN. By filtering for critical stop-gained, stop-lost, and start-lost SNPs, we narrowed down the number of candidates from over 6 million to 342 (Figure 2c, Data S1). Next, we investigated whether the genes harboring these critical SNPs are associated with plant regeneration. Specifically, we identified Arabidopsis homologs of the pepper genes containing these SNPs and compared them with previously reported regeneration-related genes (Data S2). Notably, this comparative functional analysis highlighted two genes, *PAT1* and *LONELY GUY 4* (*LOG4*). In TGN, the critical SNP in *PAT1* converts the stop codon (TGA) to Arginine (CGA), while the SNP in *LOG4* introduces a premature stop codon (TAA) in place of Lysine (AAA). However, we excluded *LOG4* from our candidates because it encodes a cytokinin riboside 5’-monophosphate phosphoribohydrolase involved in active cytokinin biogenesis (Chickarmane et al. 2012), and the SIM already contains a sufficient amount of zeatin. Therefore, we finally selected a single critical SNP occurring in the pepper *PAT1* (*CaPAT1; CA01g26680*) gene as a strong candidate for the high shoot regeneration capacity in TGN.

### TGN Harbors Full-Length CaPAT1

Since the stop-lost SNP was identified in *CaPAT1* of TGN, we analyzed the gene structure using the Jbrowse genome browser. In both N and the reference cultivar CM334, the nucleotide T478 forms a stop codon (Figure 2d). Notably, the start codon of an adjacent gene, *CA01g26690*, is located only 36 bp downstream of the *CaPAT1* stop codon (Figure S1). In TGN, a T-to-C substitution at this position converts the stop codon into an Arg codon. Because the 36-bp intergenic gap maintains the reading frame of *CA01g26690*, this substitution merges *CaPAT1* and *CA01g26690* into a single, large-sized gene (Figure 2d). Accordingly, we designated the *CaPAT1* version in N as *CaPAT1^N^*, and the merged gene in TGN as *CaPAT1^TGN^*. Protein domain analysis revealed that both CaPAT1^N^ and CA01g26690 harbor only truncated GRAS domain, whereas CaPAT1^TGN^ possesses a complete, intact GRAS domain. Furthermore, protein structure prediction using AlphaFold3 showed that the merged protein CaPAT1^T478C^ exhibits higher structural similarity to the Arabidopsis PAT1 (AtPAT1) than either CaPAT1^N^ or CA01g26690 alone (Figure 2e). These results indicate that the T478C SNP triggers a stop-to-Arg transition, inducing a gene fusion event that produces a full-length CaPAT1 with an intact GRAS domain in TGN.

### The Full-Length CaPAT1 Exhibits High Dimerization Capacity

To examine the expression patterns of CaPAT1 in N and TGN during shoot regeneration, we performed RT-qPCR analysis using *CaPAT1^N^*-specific and *CA01g26690*-specific primers. The *CaPAT1^N^*-specific primers targeted both *CaPAT1^N^* in N and *CaPAT1^TGN^*in TGN. However, no statistically significant differences in expression levels were observed during shoot regeneration (Figure 3a, upper panel). The *CA01g26690*-specific primers amplified the native *CA01g26690* in N, whereas they targeted the *CaPAT1^TGN^* resulting from the gene fusion in TGN. Similarly, no significant changes in expression were detected using these primers (Figure 3a, lower panel). These findings suggest that the transcript levels of *CaPAT1* are comparable between TGN and N throughout the shoot regeneration process.

**FIGURE 3.**
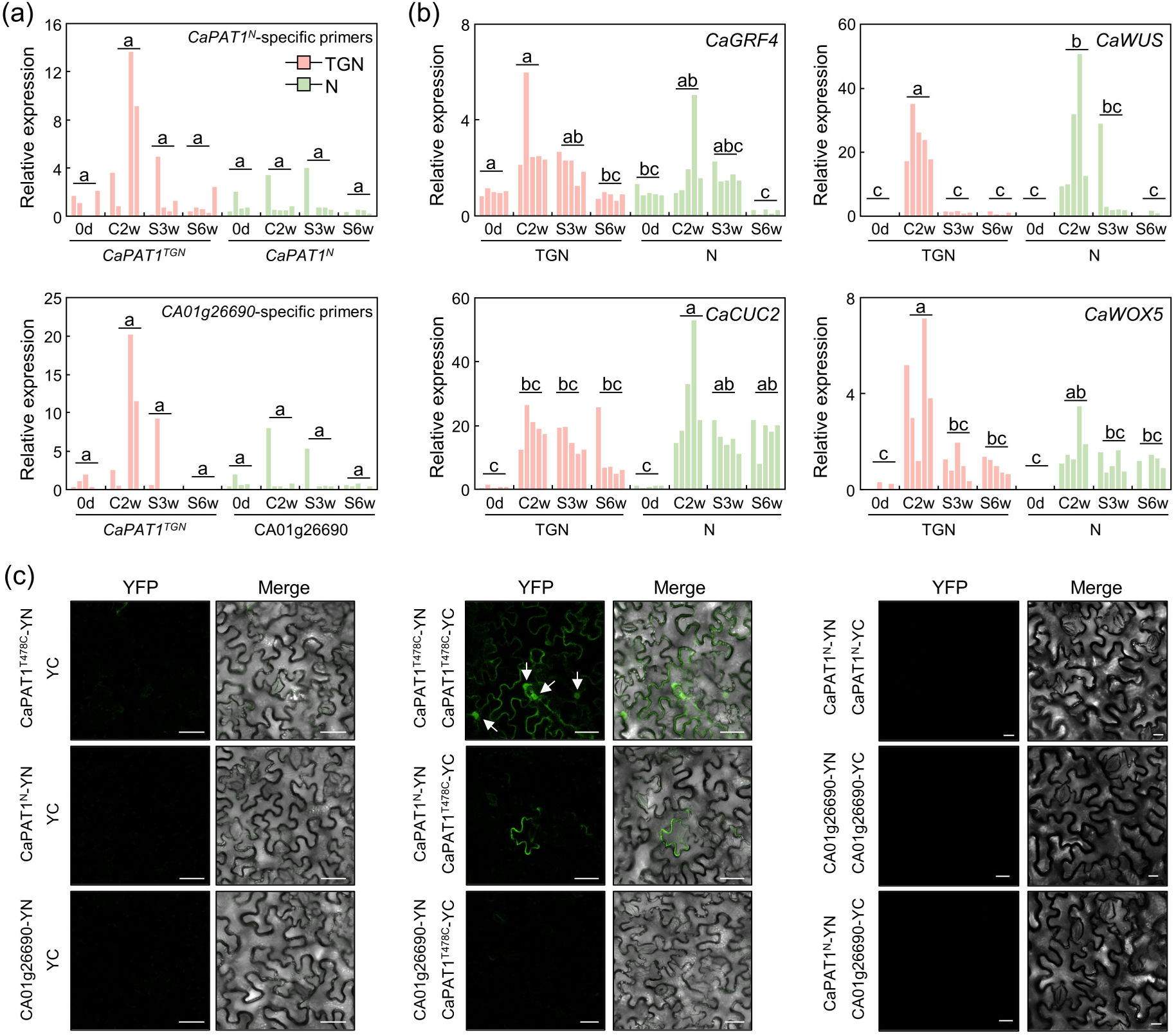
Full-length CaPAT1 is required for homodimerization. (a) Expression of *CaPAT1* and *CA01g26690* during shoot regeneration. Gene expression levels for each biological replicate are displayed. Primers specifically designed for *CaPAT1^N^* were used to analyze the expression of both *CaPAT1^N^* and *CaPAT1^TGN^* (upper panel), while those designed for *CA01g26690* were used to analyze both *CA01g26690* and *CaPAT1^TGN^* (lower panel). Different letters indicate statistically significant difference (Tukey’s test, *P* < 0.05; *n* = 5; 40 explants for each replicate). C, CIM. (b) Expression of pepper homologs of Arabidopsis regeneration-related genes. Gene expression levels for each biological replicates are displayed. Different letters indicate statistically significant difference (Tukey’s test, *P* < 0.05; *n* = 5; 40 explants for each replicate). (c) BiFC analysis of the interaction between different forms of CaPAT1. *Nicotiana benthamiana* leaves were used for transient expression analysis. YN and YC indicate the N-terminal and C-terminal halves of YFP, respectively. YFP fluorescence and bright-field images were overlapped for Merge images. Scale bars, 20 μm.

Next, we examined the expression of genes known to play key roles in inducing shoot development [*GROWTH-REGULATING FACTOR 4* (*GRF4*) and *WUS*] and pluripotency acquisition (*CUC2* and *WOX5*) during shoot regeneration (Aida et al. 1999; Pulianmackal et al. 2014; Lee et al. 2022; Debernardi et al. 2020). Pepper homologs of these four Arabidopsis genes were analyzed. While *CaWUS*, *CaCUC2*, and *CaWOX5* exhibited increased expression during shoot regeneration in both TGN and N, none of these genes showed significantly lower expression in N compared to TGN (Figure 3b). These results suggest that the transcriptional regulation of these key regeneration-related genes is not the primary cause for the low shoot regeneration capacity observed in N.

GRAS domain proteins are known to function through dimerization, a process critical for their transcription-regulatory activities (Bisht et al. 2023; Heyman et al. 2016). Given our finding that the T478C SNP in TGN induces a structural change in CaPAT1 via gene fusion, we investigated the dimerization capacities of CaPAT1^T478C^, CaPAT1^N^, and CA01g26690. In a bimolecular fluorescence complementation (BiFC) assay, CaPAT1^T478C^ homodimerization was strongly detected in both the plasma membrane and the nucleus. In contrast, CaPAT1^N^-CaPAT1^T478C^ dimers were only rarely observed and failed to localize to the nucleus (Figure 3c). Notably, no dimer formation was detected for any other combinations, including CA01g26690-CaPAT1^T478C^, CaPAT1^N^ homodimers, CA01g26690 homodimers, or CaPAT1^N^-CA01g26690 pairs. These results indicate that the full-length CaPAT1 is essential for robust dimerization and proper nuclear localization.

### CaPAT1^T478C^ Improv es Shoot Regeneration and Transformation

To verify whether the T478C SNP in *CaPAT1* determines shoot regeneration efficiency, we transformed the low-efficiency cultivar N with overexpression vectors for both *CaPAT1^N^* and *CaPAT1^T478C^*. Analysis of shoot regeneration frequency in the transformed T0 generation revealed that overexpression of *CaPAT1^N^* did not significantly alter regeneration efficiency compared to the empty vector (EV) control (Figure 4a). In contrast, ectopic expression of *CaPAT1^T478C^* significantly enhanced shoot regeneration frequency (Figure 4b,c). Notably, the efficiency of generating transgenic shoots was also significantly improved by *CaPAT1^T478C^* transformation. We obtained total 12 transgenic shoots from 156 cultured explants, whereas EV transformation yielded only a single transgenic shoot from 148 explants (Figure 4d). These observations suggest that the T478C SNP in *CaPAT1* is closely related to improving shoot regeneration efficiency in pepper.

**FIGURE 4.**
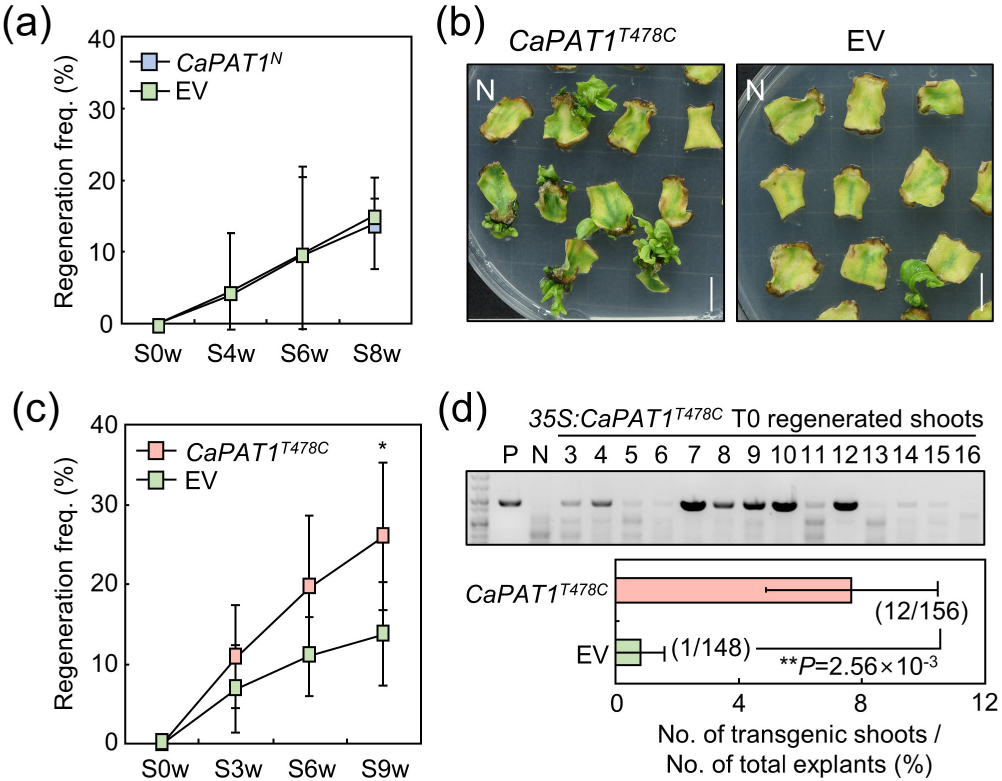
Ectopic expression of *CaPAT1^T478C^* enhances shoot regeneration capacity. (a) Regeneration frequency following *CaPAT^N^* overexpression. Leaf explants of cultivar N were transformed with either an EV or the *CaPAT1^N^*-overexpressing vector. Shoot regeneration was analyzed in the transformed T0 generation. No statistically significant differences were detected by Student’s *t*-test (*n* = 4; 113 explants for each replicate). (b, c) Effects of ectopic *CaPAT1^T478C^* expression on shoot regeneration. Either EV or the *CaPAT1^T478C^*-overexpressing vector was transformed into leaf explants of N. Representative images are displayed (b). Regeneration frequency was analyzed at the indicated time points (c). Statistical significance was determined using Student’s *t*-test (\**P* < 0.05; *n* = 6; 85 explants for each replicate). Scale bars, 1cm. (d) Efficiency of transgenic shoot production via *CaPAT^T478C^* overexpression. Regenerated shoots in (c) were analyzed for transgene integration. Statistical significance was determined using Student’s *t*-test (*n* = 5). The total number of regenerated transgenic shoots and the total number of explants across all independent experiments are indicated in the graph. For line and bar graphs, whiskers indicate ±SD.

We next analyzed the presence of T478C SNP across various pepper genotypes. Most cultivars, including CM334, Dempsey, and several commercial varieties, harbored the N-type *CaPAT1* sequence (T478). However, two inbred lines, C15 and P915, possessed the T478C SNP, similar to TGN (Figure 5a). To investigate whether *CaPAT1^T478C^* overexpression remains effective in genotypes with different *CaPAT1* backgrounds, we transformed C15 with the *CaPAT1^T478C^* overexpression vector. Although C15 already harbors the T478C SNP, ectopic expression of *CaPAT1^T478C^* further enhanced its shoot regeneration efficiency (Figure 5b,c). Given that C15 possesses a high innate capacity for shoot growth and rooting, we evaluated transformation efficiency in pot-grown plants derived from the regenerated shoots (Figure 5d). Notably, transformation efficiency was significantly elevated, yielding total 30 transgenic plants from 74 cultured explants, compared to only a single transgenic plant from the EV control (Figure 5e). These results suggest that the expression level of *CaPAT1* is important for high shoot regeneration capacity.

**FIGURE 5.**
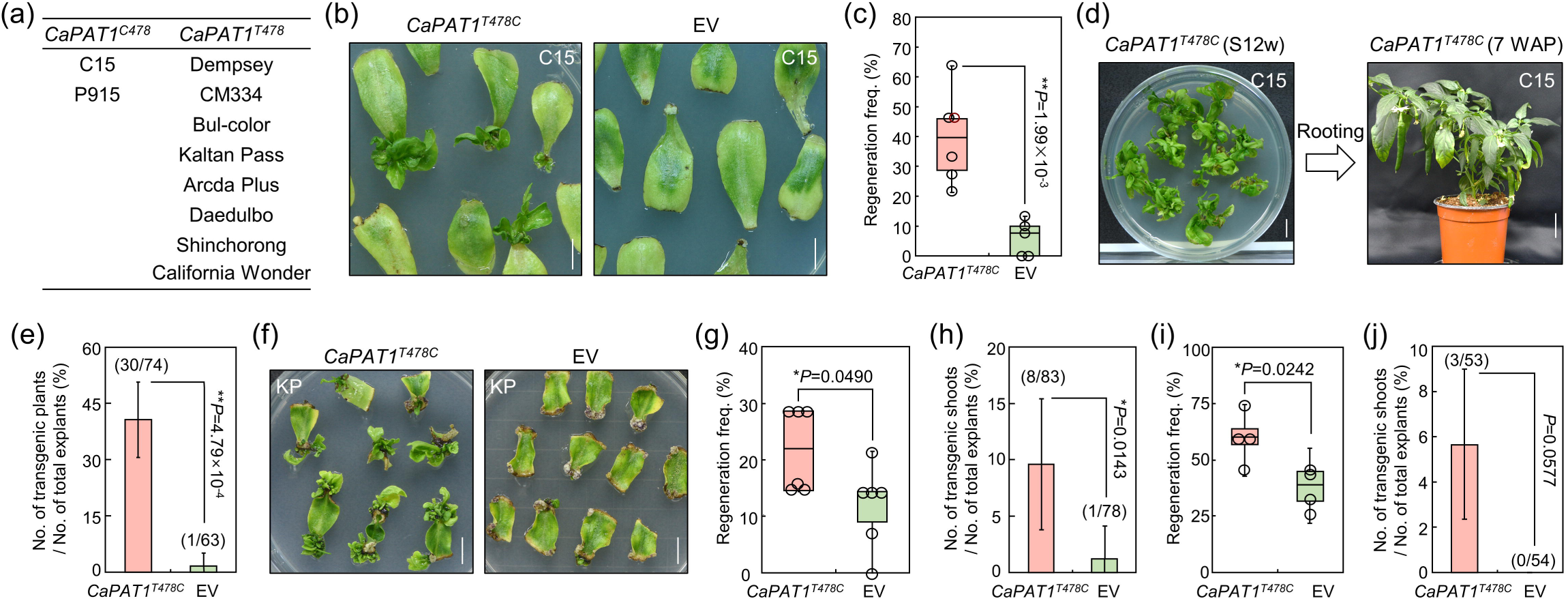
Ectopic expression of *CaPAT1^T478C^* enhances shoot regeneration and transformation efficiency across multiple genotypes. (a) *CaPAT1* types in multiple pepper cultivars. The *CaPAT1* locus in diverse pepper genotypes was analyzed via Sanger sequencing. (b, c) Effects of ectopic *CaPAT1^T478C^* expression on shoot regeneration in C15. Shoot regeneration in the transformed T0 generation was analyzed. Representative images are displayed (b). Regeneration frequency was determined at S10w (c). Statistical significance was determined using Student’s *t*-test (*n* = 5; 12 explants for each replicate). Scale bars, 1cm. (d, e) Transformation efficiency following *CaPAT1^T478C^* overexpression in C15. Regenerated shoots obtained in (c) were transferred to hormone-free medium for rooting (d). After rooting, plants were transferred to soil. Leaves from soil-grown plants were harvested to analyze transgene integration. Efficiency of generating transgenic plants was calculated (e). Statistical significance was determined using Student’s *t*-test (*n* = 5). Scale bars, 1 cm. (f–h) Effects of ectopic *CaPAT1^T478C^* expression in Kaltan Pass (KP). Representative images are displayed (f). Regeneration frequency was analyzed at S8w (g). Statistical significance was determined using Student’s *t*-test (*n* = 6; 13 explants for each replicate). Efficiency of generating transgenic shoots was calculated (h). Statistical significance was determined using Student’s *t*-test (*n* = 6). Scale bars, 1 cm. (i, j) Effects of ectopic *CaPAT1^T478C^* expression in California Wonder. Regeneration frequency was analyzed at S8w (i). Statistical significance was determined using Student’s *t*-test (*n* = 4; 13 explants for each replicate). Efficiency of generating transgenic shoots was calculated (j). Statistical significance was determined using Student’s *t*-test (*n* = 4). For bar graphs, whiskers indicate ±SD. For (e), (h), and (j), the total number of obtained transgenic shoots and explants across all independent experiments are indicated in the graph.

We also validated the effects of *CaPAT1^T478C^* overexpression in the commercial cultivar Kaltan Pass. Consistent with our findings, *CaPAT1^T478C^* overexpression significantly increased both shoot regeneration frequency and transformation efficiency (Figure 5f–h). We obtained total 8 transgenic shoots from 83 cultured explants following *CaPAT1^T478C^*overexpression, whereas EV transformation yielded only one transgenic plant from 78 explants (Figure 5h). Next, we examined the efficacy of *CaPAT1^T478C^* in bell pepper, California Wonder. Transformation with the *CaPAT1^T478C^*overexpression vector led to a significant increase in shoot regeneration efficiency (Figure 5i). Notably, while no transgenic shoots were obtained from the EV control, total three transgenic shoots were obtained from 53 explants transformed with *CaPAT1^T478C^* (Figure 5j). Collectively, these results demonstrate that the overexpression of *CaPAT1^T478C^* serves as a robust tool to overcome recalcitrance in shoot regeneration and transformation across diverse pepper genotypes.

## Discussion

In this study, we identified that the T478C SNP in CaPAT1 is a key determinant underlying shoot regeneration capacity in pepper by comparing a high-efficiency genotype (TGN) with a low-efficiency genotype (N). Importantly, overexpression of the *CaPAT1^T478C^* significantly improved both shoot regeneration capacity and transformation efficiency across diverse pepper genotypes. These results suggest that restoring full-length functional domains of CaPAT1 provides a practical and broadly applicable strategy to overcome regeneration barriers in pepper. Efficient transformation is a prerequisite for genome editing. Therefore, these results suggest that manipulation of *CaPAT1* provides a practical and broadly applicable strategy not only to overcome regeneration barriers in pepper but also to facilitate the efficient production of gene-edited plants. Given the high conservation of GRAS family proteins across diverse plant species (Sun et al. 2012; Wang et al. 2018), this mechanism likely extends beyond pepper. Therefore, identifying critical SNPs such as stop-lost, stop-gained, or start-lost polymorphisms within *PAT1* or homologous genes encoding GRAS-domain proteins and subsequently engineering these SNPs represents a promising strategy to overcome recalcitrance in shoot regeneration across a broad range of crops.

In Arabidopsis, *PAT1* has been reported to function as a positive regulator of phytochrome A-mediated light signaling and to interact with key transcriptional regulators (Bolle et al. 2004; Torres-Galea et al. 2013). Beyond its role in light signaling, *PAT1* is also involved in plant regeneration processes. For instance, PAT1 forms a heterodimer with ERF115 to enhance the regeneration of root stem cells (Heyman et al. 2016). Overexpression of both *PAT1* and *ERF115* triggers spontaneous callus formation and the upregulation of genes such as *PLT3*, *PLT5* and *PLT7*, which are central genes conferring pluripotency for shoot regeneration in callus (Kareem et al. 2015). Notably, a recent study showed that *PAT1* is necessary for efficient grafting in both Arabidopsis and Norway spruce (*Picea abies*) (Feng et al. 2024). Arabidopsis mutants deficient in both *PAT1* and its close homolog *SCL5* exhibited disrupted grafting efficiency. These findings suggest that *PAT1* may also be involved in shoot regeneration, although this function has not been previously examined. In this study, we first demonstrated that *PAT1* is a key gene determining the efficiency of shoot regeneration in pepper. Further research on the signaling pathways of *CaPAT1* utilizing single-cell RNA sequencing and subsequent analysis of the transcriptome in *CaPAT1*-expressing cells will provide insights into how CaPAT1 facilitates shoot regeneration in callus cells.

In this study, we observed that full-length CaPAT1 efficiently forms homodimers, whereas truncated CaPAT1s exhibit low dimerization activities (Figure 3c). AlphaFold-based protein structure prediction revealed that full-length CaPAT1^T478C^ harbors both an α-helical cap and an α/β subdomain. In contrast, CaPAT1^N^ possesses a truncated α/β domain, and CA01g26690 lacks both the α-helical cap and a partial α/β subdomain (Figure 2e). These results suggest that the α/β subdomain plays a major role in the dimerization of CaPAT1. This is consistent with previous findings indicating that a hydrophobic groove within the α/β subdomain is critical for protein-protein interactions (Hakoshima et al. 2018). Because GRAS proteins often form heterodimeric complexes to execute their regulatory functions (Helariutta et al. 2000; Sabatini et al. 2003), the full-length GRAS domain in CaPAT1 may be necessary for heterodimerization with other GRAS proteins, such as SCL5 and SCL21, as well as for homodimerization. Screening for protein interactions among CaPAT1^T478C^, CaPAT1^N^, CA01g26690, and other CaPAT1 homologs in pepper will be helpful to elucidate the molecular functions of CaPAT1 during shoot regeneration.

## Materials and Methods

### Plant Materials and Seed Germination

The N (cat. No: 02-0004-2013-56) and Bul-color (cat. No: 00-1997-10-01) seeds were purchased from Farmhannong. The Kaltan Pass seeds were purchased from Nongwoobio (cat. No: 10-1998-10-01). The Arcda Plus seeds were purchased from Hyundai Seed (cat. No: 10-2001-10-16). The Daedulbo seeds were purchased from Takii Seed (cat. No: 10-2003-10-22). The Shinchorong seeds were purchased from Sakata Korea (cat. No: 10-2004-10-26). The California Wonder seeds were purchased from Premiumseeds (cat. No: 80249CIN0EE). The TGN seeds were provided as a gift by Chonnam National University. The C15, CM334, Dempsey, and P915 seeds were a gift from Nongwoo Seed. The pepper seeds from each genotype were sterilized by incubating them in 70% EtOH for 30 sec, followed by shaking them in a 5% NaOCl solution containing 0.01% Tween 20 for 10 min. Subsequently, the seeds were rinsed with sterile distilled water eight times and then dried on sterilized filter paper. After sterilization, the seeds were germinated on MS-agar medium [2.15 g L^−1^ MS basal salt mixture (Duchefa, cat. No: M0222.0050), 1.5% sucrose, and 0.6% phyto agar (Duchefa, cat. No: P1003) with a pH 5.8]. The plates were incubated in the dark for 7 days; then, they were transferred to long-day conditions (16-hour light and 8-hour dark) at 24°C.

### Shoot Regeneration from Pepper Explants

To induce shoot regeneration, cotyledons of 11-day-old seedlings were excised into 0.5 cm pieces and incubated onto CIM [MS-agar medium containing 2.0 mg L^−1^ of zeatin and 0.2 mg L^−1^ of indole-3-acetic acid (IAA)] at 24°C under dark conditions. After 4 weeks, they were transferred to SIM (MS-agar medium containing 2.0 mg L^−1^ of zeatin and 0.05 mg L^−1^ of IAA) at 24°C under long-day conditions. Explants were transferred to the fresh SIM every two weeks. The regeneration frequency was determined by dividing the number of explants showing regenerated shoots by the total number of explants.

### Whole Genome-Sequencing and Single Nucleotide Polymorphism Analysis

Genomic DNA was extracted from leaves of NK and TGN using the HiYield^TM^ Genomic DNA Mini Kit (Plant) (RBC, cat. No: TGP100). A WGS library was constructed using the QIAseq FX Library CDI kit (Qiagen), and next-generation sequencing of the two pepper lines was performed using the Illumina platform. CM334 genome was used as a reference (Kim et al. 2014). SNPs were classified as homozygous SNPs if the mapped reads at the given position were identical in more than 90% of the cases, and as heterozygous SNPs if 40% to less than 60% of the reads were different. All other cases were classified as etc.

### Analysis of Protein Structure

Protein structure prediction was performed using AlphaFold3. PyMOL software was used to display protein structures. To determine structural similarity between Arabidopsis and pepper PAT1s, TM scores were calculated using the TM-align tool (https://zhanggroup.org/TM-align/).

### BiFC Assay

To visualize protein-protein interactions in plant cells, the BiFC vectors *pSPYNE* and *pSPYCE,* (for split YFP N-terminal/C-terminal fragment expression) were used. The coding sequence of *CaPAT1^T478C^*, *CaPAT1^N^* and *CA01g26690* were cloned into the *pSPYCE-35S* and *pSPYCE-35S* vectors. The resulting constructs were delivered into leaf cells of *Nicotiana benthamiana* via *Agrobacterium tumefaciens* infiltration. For transient expression, *Agrobacterium* cells containing *pSPYCE-35S* and *pSPYCE-35S* were harvested by centrifugation, resuspended, and diluted in infiltration buffer (10 mM MES, 10 mM MgSO_4_, 200 μM acetosyringone, at pH 5.7) to an OD_600_ = 1.0. Equal volumes of *Agrobacterium* suspensions carrying the corresponding constructs were mixed and syringe-infiltrated into the abaxial side of leaves from 4-week-old *N. benthamiana* plant leaves. Leaf tissues were harvested on the 4th days post-infiltration and observed using a confocal laser scanning microscope. YFP fluorescence was excited at 488 nm, and images were analyzed using ZEN 2.5 LITE software.

### RNA Extraction and RT-qPCR Analysis

A portion of the explants located in proximity to the wound site and containing calli was excised for extracting total RNA using the RNeasy Plant Mini kit (Qiagen, cat. No: 74904). For RT-qPCR, 1 μg of total RNA was used to synthesize complementary DNA (cDNA) using TOPscript cDNA Synthesis kit (Enzynomics, cat. No: EZ005M). RT-qPCR was performed using TOPreal SYBR Green qPCR PreMIX (Enzynomics, cat. No: RT500M) in CFX Connect real-time PCR detection system (Bio-Rad, cat. No: 1855201). Primers used for RT-qPCR are listed in Table S1. The *CaUBQ1* was used as a reference gene.

### *Agrobacterium*-Mediate Transformation

The *CaPAT1^N^* or *CaPAT1^T478C^* sequences were inserted into the pK2GW7 vector using the gateway cloning method. The *CaPAT1^N^*or *CaPAT1^T478C^*-overexpressing vector was introduced into *Agrobacterium* strain GV3101. *Agrobacterium* cells were cultured in YEP medium [10 g L^−1^ peptone (Gibco, cat. No: 211677), 10 g L^−1^ yeast extract (Gibco, cat. No: 211929), and 5 g L^−1^ NaCl (Samchun, cat. No: S0484)] until reaching OD_600_ = 0.5. The cells were then harvested by centrifugation and resuspended in 40 ml of MS-liquid medium containing 100 μM acetosyringone without sucrose. Cotyledon explants from the all genotypes except C15 cultivar were induced to from callus in CIM for 2 weeks, then co-cultivated in *Agrobacterium*-resuspended MS medium for 30 min with shaking every 5 min. Subsequently, the explants were placed onto sterile filter paper to remove excess liquid medium and incubated in darkness on co-cultivated medium (MS-agar medium containing 100 μM acetosyringone) for 4 days. The infected explants were washed by incubating them in 100 mg L^−1^ cefotaxime solution for 5 min. Subsequently, the explants were dried on sterilized filter paper and then cultured on SIM containing 3.0 mg L^−1^ 6-benzylaminopurine, 0.05 mg L^−1^ IAA, 2 mg L^−1^ AgNO_3_, 500 mg L^−1^ cefotaxime, and appropriate selection markers under long-day conditions, with the medium changed every 2 weeks until shoots regenerated from the callus.

Cotyledon explants of the C15 were co-cultivated in *Agrobacterium*-resuspended MS medium for 30 min with shaking every 5 min. Subsequently, the explants were placed onto sterile filter paper to remove excess liquid medium and incubated in the dark co-cultivated medium for 2 days. The explants were then cultured on CIM containing 500 mg L^−1^ cefotaxime, and appropriate selection markers under dark conditions, with the medium changed every 2 weeks for a total of 4 weeks. After CIM incubation, the explants were transferred to SIM containing 500 mg L^−1^ cefotaxime, and appropriate selection markers under long-day conditions, with the medium changed every 2 weeks until shoots regenerated from the callus.

### Genomic DNA Extraction and PCR Analysis

Genomic DNA was extracted from pepper leaf using HiYield^TM^ Genomic DNA Mini Kit (RBC, cat. No: YGP100). To analyze the *CaPAT1* locus in multiple pepper cultivars, PCR was performed using 2 × Pfu PCR Master Mix (BIOFACT, cat. No: PD301-50h). The PCR conditions were initial denaturation at 95°C for 2 min, followed by 35 cycles of 95°C for 30 sec, 60°C for 30 sec and 72°C for 1 min, with a final extension at 72°C for 5 min. To analyze the integration of the transgene, PCR was performed using 2 × TOP simple^TM^ PCR DyeMIX-Tenuto (Enzynomics, cat. No: P510). The PCR conditions included the initial denaturation at 95°C for 2 min, followed by 35cycles of 95°C for 30 sec, 58°C for 30 sec and 72°C for 1 min, with a final extension at 72°C for 5 min. Primers used for PCR are listed in Table S1.

### Statistical Analysis

All statistical methods and the number of biological or technical replicates in each assay are described in the figure legends. Student’s *t*-test was performed using Microsoft Excel 2016. One-way ANOVA followed by Tukey’s HSD test was conducted using Rstudio.

## Supporting information

Supporing Figure

## Author Contributions

S.J.P., H.S.K., and H.J.L. conceived the research and designed the experiments. S.J.P. performed all experiments and conducted data analysis. J.H.Y. assisted with the experimental procedures. S.J.P. and H.J.L. wrote the manuscript, and H.S.K. provided critical revisions and edited the manuscript.

## Data Availability Statement

Whole genome sequencing raw data for TGN and N are deposited in K-BDS (https://kbds.re.kr/). Accession number is KAP242334.

## References

Aida, M., T. Ishida, and M. Tasaka. 1999. “Shoot Apical Meristem and Cotyledon Formation during Arabidopsis Embryogenesis: Interaction among the CUP- SHAPED COTYLEDON and SHOOT MERISTEMLESS Genes.” Development 126:1563–1570.

Bisht, A., T. Eekhout, B. Canher, et al. 2023. “PAT1-type GRAS-domain Proteins Control Regeneration by Activating DOF3.4 to Drive Cell Proliferation in Arabidopsis Roots.” Plant Cell 35, 1513–1531.

Bolle, C. 2004. “The Role of GRAS Proteins in Plant Signal Transduction and Development.” Planta 218, 683–692.

Bustillo-Avendaño, E., S. Ibáñez, O. Sanz, et al. 2017. “Regulation of Hormonal Control, Cell Reprogramming and Patterning during *de novo* Root Organogenesis.” Plant Physiology 176, 1709–1727.

Chickarmane, V.-S., S.-P. Gordon, P.-T. Tarr, M.-G. Heisler, and E.-M. Meyerowitz. 2012. “Cytokinin Signaling as a Positional Cue for Patterning the Apical–basal Axis of the Growing Arabidopsis Shoot Meristem.” Proceedings of the National Academy of Sciences of the United States of America 109, 4002–4007.

Christianson, M.-L., and D.-A. Warnick. 1983. “Competence and Determination in the Process of *in vitro* Shoot Organogenesis.” Developmental Biology 95, 288–293.

Debernardi, J.-M., D.-M. Tricoli, M.-F. Ercoli, et al. 2020. “A GRF-GIF Chimeric Protein Improves the Regeneration Efficiency of Transgenic Plants.” Nature Biotechnology 38, 1274–1279.

Feng, M., A. Zhang, V. Nguyen, et al. 2024. “A Conserved Graft Formation Process in Norway Spruce and Arabidopsis Identifies the PAT Gene Family as Central Regulators of Wound Healing.” Nature Plants 10,53–65.

Gammoudi, N., T.-S. Pedro, A. Ferchichi, and C. Gisbert. 2018. “Improvement of Regeneration in Pepper: a Recalcitrant Species.” In Vitro Cellular Developmental Biology-Plant 54, 145–153.

Gordon, S.-P., V.-S. Chickarmane, C. Ohno, and E.-M. Meyerowitz. 2009. “Multiple Feedback Loops Through Cytokinin Signaling Control Stem Cell Number within the Arabidopsis Shoot Meristem.” Proceedings of the National Academy of Sciences of the United States of America 106, 16529–16534.

Gubis, J., Z. Lajchova, J. Farago, and Z. Jurekova. 2004. “Effect of Growth Regulators on Shoot Induction and Plant Regeneration in Tomato (*Lycopersicon esculentum* Mill.).” Biologia 59, 405–408.

Hakoshima, T. 2018. “Structural Basis of the Specific Interactions of GRAS Family Proteins.” Federation of European Biochemical Societies 4, 489–501.

Helariutta, Y., H. Fukaki, J. Wysocka-Diller, et al. 2000. “The *SHORT-ROOT* Gene Controls Radial Patterning of the Arabidopsis Root through Radial Signaling.” Cell 101, 555–567.

Hernandez-Perez, T., M.-D. Gomez-Garcia, M.-E. Valverde and O. Paredes-Lopez. 2020. “*Capsicum annuum* (hot pepper): An ancient Latin-American Crop with Outstanding Bioactive Compounds and Nutraceutical Potential. A review.” Comprehensive Reviews in Food Science and Food Safety 19, 2972–2993.

Heszky, L.-E., D.-Q. Binh, E. Kiss, and G. Gyulai. 1989. “Increase of Green Plant Regeneration Efficiency by Callus Selection in Puccinellia limosa (Schur.) Holmbg.” Plant Cell Reports 3, 174–177.

Heyman, J., T. Cools, B. Canher, et al. 2016. “The Heterodimeric Transcription Factor Complex ERF115-PAT1 Grants Regeneration Competence.” Nature Plants 2, 16165.

Higuchi, M., M.-S. Pischke, A.-P. Mähönen, et al. 2004. “*In planta* Functions of the Arabidopsis Cytokinin Receptor Family.” Proceedings of the National Academy of Sciences of the United States of America 101, 8821–8826.

Ikeuchi, M., D.-S. Favero, Y. Sakamoto, et al. 2019. “Molecular Mechanisms of Plant Regeneration.” Annual Review of Plant Biology 70, 377–406.

Kareem, A., K. Durgaprasad, K. Sugimoto, et al. 2015. “*PLETHORA* Genes Control Regeneration by a Two-step Mechanism.” Current Biology 25, 1017–1030.

Kim, M.-S., Y.-J. Han, T. Tripathi, et al. 2023. “Comparison of Regeneration Conditions in Seven Pepper (*Capsicum annuum* L.) Varieties.” Korean Journal of Plant Resources 36, 527–539.

Kim, S., M. Park, S.-I. Yeom, et al. 2014. “Genome Sequence of the Hot Pepper Provides Insights into the Evolution of Pungency in Capsicum Species.” Nature Genetics 3, 270–278.

Ko, M., J.-H. Cho, H.-H. Seo, et al. 2016. “Constitutive Expression of a Fungus-inducible Carboxylesterase Improves Disease Resistance in Transgenic Pepper Plants.” Planta 244, 379–392.

Kothari, S.-L, A. Joshi, S. Kachhwaha, and N. Ochoa-Alejo. 2010. “Chilli Peppers - A review on Tissue Culture and Transgenesis.” Biotechnology Advances 28, 35–48.

Lee, K., J.-H. Kim, O.-S. Park, Y.-J. Jung, and P.-J. Seo. 2022. “Ectopic Expression of *WOX5* Promotes Cytokinin Signaling and *de novo* Shoot Regeneration.” Plant Cell Reports 41, 2415–2422.

Lee, K., and P.-J. Seo. 2018. “Dynamic Epigenetic Changes during Plant Regeneration.” Trends in Plant Science 23, 235–247.

Torres-Galea, P., B. Hirtreiter, and C. Bolle. 2013. “Two GRAS Protein, SCARECROW-LIKE21 and PHYTOCHROME A SIGNAL TRANSDUCTION1, Function Cooperatively in Phytochrome A Signal Transduction.” Plant Physiology 161, 291–304.

Liu, J., L. Sheng, Y. Xu, et al. 2014. “*WOX 11* and *12* are Involved in the First-step Cell Fate Transition during *de novo* Root Organogenesis in Arabidopsis.” Plant Cell 26, 1081–1093.

Mazumdar, P., A. Basu, A. Paul, C. Mahanta, and L. Sahoo. 2010. “Age and Orientation of the Cotyledonary Leaf Explants Determine the Efficiency of *de novo* Plant Regeneration and *Agrobacterium tumefaciens*-mediated Transformation in *Jatropha curcas* L.” South African Journal of Botany 76, 337–344.

Meng, W.-J., Z.-J. Cheng, Y.L. Sang, et al. 2017. “Type-B ARABIDOPSIS RESPONSE REGULATORs Specify the Shoot Stem Cell Niche by Dual Regulation of *WUSCHEL*.” Plant Cell 29, 1357–1372.

Naeem, B., S. Shams, L. Ma, et al. 2025. “An Optimized Regeneration Protocol for Chili Peppers (*Capsicum annuum* L.) through Genotype-specific Explant and Growth Regulator Combinations.” Seed Biology 4, e012.

Park, J.-S., K.-H. Park, S.-J. Park, et al. 2023. “WUSCHEL Controls Genotype-Dependent shoot regeneration capacity in potato.” Plant Physiology 193, 661–676.

Park, S.-I., H.-B. Kim, H.-J. Jeon, and H. Kim, H. 2021. “*Agrobacterium*-Mediated *Capsicum annuum* Gene Editing in Two Cultivars, Hot Pepper CM334 and Bell Pepper Dempsey.” International Journal of Molecular Science 8, 3921.

Pulianmackal, A.-J., A.V. Kareem, K. Durgaprasad, Z. B. Trivedi, and K. Prasad. 2014. “Competence and Regulatory Interactions during Regeneration in Plants.” Frontiers in Plant Science 5, 142.

Radhakrishnan, D., A. Kareem, K. Durgaprasad, et al. 2018. “Shoot Regeneration: a Journey from Acquisition of Competence to Completion.” Current Opinion in Plant Biology 41, 23–31.

Rajametov, S. N., E. Y. Yang, M. C. Cho, et al. 2021. “Heat-Tolerant Hot Pepper Exhibits Constant Photosynthesis via Increased Transpiration Rate, High Proline Content and Fast Recovery in Heat Stress Condition.” Scientific Reports 11,14328.

Sabatini, S., R. Heidstra, M. Wildwater, and B. Scheres. 2003. “SCARECROW is Involved in Positioning the Stem Cell Niche in the Arabidopsis Root Meristem.” Genes and Development 17, 354–358.

Seo, H.-H., S. Park, S. Park, et al. 2014. “Overexpression of a Defensin Enhances Resistance to a Fruit-Specific Anthracnose Fungus in Pepper.” Public Library of Science One 9, e97936.

Shams, S., B. Naeem, L. Ma, et al. 2024. “Developing an Optimized Protocol for Regeneration and Transformation in Pepper.” Genes 15, 1018.

Shin, M. J., S. S. Ku, S. -J. Park, S. U. Park, S. R. Min, H. -S. Kim, and H. -J. Lee. 2026. “Suppressing microRNA396 activity enhances regeneration efficiency of viable shoots from cotyledon explants in pepper.” Plant Biotechnology Reports 20, 15.

Suica-Bunghez, I.-R., and R.-M. Ion. 2020. “Characterization, Phytochemical and Antioxidant Activity of Three Types of Hot Pepper (*Capsicum Annuum* L.).” Journal of Science and Arts 51, 443–450.

Sun, X., W. T. Jones, and E. H. A. Rikkerink. 2012. “GRAS Proteins: the Versatile Roles of Intrinsically Disordered Proteins in Plant Signalling.” Biochemical journal 442, 1–12.

Ulmasov, T., G. Hagen, and T. J. Guilfoyle. 1999. “Activation and Repression of Transcription by Auxin-Response Factors.” Proceedings of the National Academy of Sciences of the United States of America 96, 5844–5849.

Wang, Y.-X., Z.-W. Liu, Z.-J. Wu, et al. 2018. “Genome-Wide Identification and Expression Analysis of GRAS Family Transcription Factors in Tea Plant (*Camellia sinensis*).” Scientific Reports 8, 3949.

Zhang, T,-Q., H. Lian, C.-M Zhou, et al. 2017. “A Two-Step Model for *de novo* Activation of *WUSCHEL* during Plant Shoot Regeneration.” Plant Cell 29, 1073–1087.

