## Supplementary material for "A full-length GRAS domain protein underpins efficient shoot regeneration in pepper": Supporing Figure

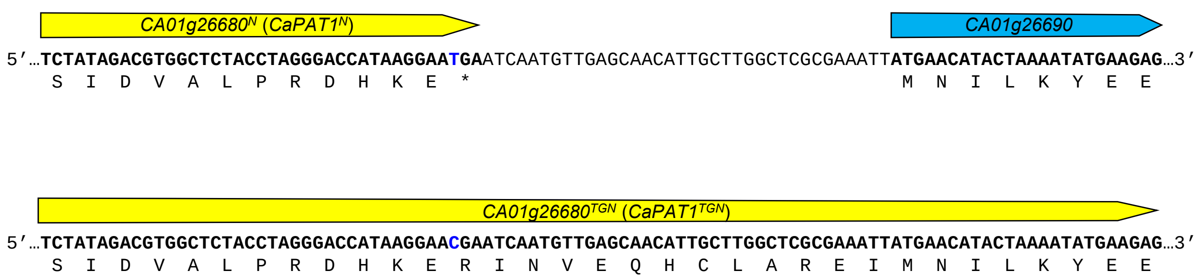


**Figure S1** | Codon analysis of *CaPAT1* in TGN and N.

Partial DNA sequences of *CaPAT1*$^{N}$ and *CA01g26690* in N and the fused *CaPAT1*$^{TGN}$ in TGN, are displayed. Codons and their corresponding amino acid sequences are indicated. Blue nucleotides represent the critical SNP that converts the stop codon in N into an Arg codon in TGN, facilitating the read-through and gene fusion.
