## Supporting Table for "A full-length GRAS domain protein underpins efficient shoot regeneration in pepper"

**Table S1** | Primers used in this study.

| **Primers** | **Sequences** | **Usage** |
| --- | --- | --- |
| CaUBQ1_qF | 5'-AAGGAAATGTGTGTCTCAAC-3' | RT-qPCR |
| CaUBQ1_qR | 5'-TCCAAATGCCAAACTTCTAG-3' | RT-qPCR |
| CaGRF4_qF | 5'-AAACCGTTCAAGAAAGCCTG-3' | RT-qPCR |
| CaGRF4-qR | 5'-TGCTTCCGTAACTCATTCCA-3' | RT-qPCR |
| CaWUS_qF | 5'-TACCACCCTGGCTTAGAAAC-3' | RT-qPCR |
| CaWUS_qR | 5'-AGACTGAGTTCAAGAGCAGC-3' | RT-qPCR |
| CaCUC2_qF | 5'-AGCACGTGTCCTGTTTCTCC-3' | RT-qPCR |
| CaCUC2_qR | 5'-GGAGAACAACGGAAGCTGGA-3' | RT-qPCR |
| CaWOX5_qF | 5'-AGACAGAAACGTCGTCGTAA-3' | RT-qPCR |
| CaWOX5_qR | 5'-GCCTGATCTGACTCTCGATT-3' | RT-qPCR |
| CA01g26680_qF | 5'-ACTACGCAGAGTGGTATCTG-3' | RT-qPCR |
| CA01g26680_qR | 5'-GTAGATGGAGCTTCCCAAGG-3' | RT-qPCR |
| CA01g26690-qF | 5'-TGAGCTTCTTGAAAGGTGGA-3' | RT-qPCR |
| CA01g26690-qR | 5'-ACCAAATTGTGATTCATCGCC-3' | RT-qPCR |
| 35s pro_F | 5'-TGAGACTTTTCAACAAAGGGTAATATC-3' | Genomic DNA PCR |
| Nos_F | 5'-GAACCGCAACGTTGAAGGAGCC-3' | Genomic DNA PCR |
| NPT_F | 5'-TGCCGAGAAAGTATCCATCATG-3' | Genomic DNA PCR |
| attB1CaPAT1_F | 5'-aaaaagcaggctccATGTCAGAACTACGCAGAGT-3' | Cloning |
| attB2CaPAT1_R | 5'-agaaagctgggttCTATACTTCTGGTAGGATGA-3' | Cloning |
| attB2CaPAT1N_R | 5'-agaaagctgggttTCaTTCCTTATGGTCCCTAGG-3' | Cloning |
| pSPYNE_PAT1F_F | 5'-tggagagaacacgggggactATGTCAGAACTACGCAGAG-3' | BiFC |
| pSPYNE_PAT1F_R | 5'-aaatcaacttttgctccatcTACTTCTGGTAGGATGAC-3' | BiFC |
| pSPYNE_PAT1C_F | 5'-tggagagaacacgggggactATGAACATACTAAAATATGAAGAGG-3' | BiFC |
| pSPYNE_PAT1C_R | 5'-aaatcaacttttgctccatcTACTTCTGGTAGGATGAC-3' | BiFC |
| pSPYNE_PAT1N_R | 5'-aaatcaacttttgctccatcTCGTTCCTTATGGTCCCTAG-3' | BiFC |
| pSPYCE_PAT1C_F | 5'-tggagagaacacgggggactATGAACATACTAAAATATGAAGAGG-3' | BiFC |
| pSPYCE_PAT1C_R | 5'-gaacatcgtatgggtacatcTACTTCTGGTAGGATGAC-3' | BiFC |
